# Invisible people: Exploring how well remote-sensed datasets reveal the distribution of forest-proximate populations

**DOI:** 10.1101/2024.09.23.614649

**Authors:** Mirindra Rakotoarisoa, Julia P. G. Jones, O. Sarobidy Rakotonarivo, Manoa Rajaonarivelo, Dominik Schüßler

## Abstract

Accurate information on the location and density of people living at the forest frontier is vital for effective and equitable forest conservation. We compare the location of settlements and estimated population density from three global-scale, remote-sensed datasets (World Settlement Footprint 2015, Open Buildings, WorldPop) with a fine-scale, manually-derived dataset of 3,136 human settlements, of which 95% had fewer than 150 households. The study region is located in north-eastern Madagascar, contains three protected areas and the largest unprotected block of humid forest of the island. The Open Buildings dataset detected a much higher proportion (94%) of settlements than did World Settlement Footprint (15%). Population density from WorldPop matches poorly with that estimated from our manually-derived dataset. The accuracy of all three datasets is worse in more remote, forested areas, further away from basic infrastructure. Open Buildings appears to best reveal the distribution of low density scattered populations in forested areas. However, further testing in other climatic regions is still needed. Making good use of appropriate remote-sensed data could revolutionize the inclusion of local communities in conservation policy and practice, improve the quality of inference in conservation research, particularly in times of a planned expansion of the global protected area network.

## Introduction

Land use change is a major driver of biodiversity decline (IPBES, 2019; Jaureguiberry et al., 2022). Subsistence farming, mostly carried out by poor people living in scattered settlements along the forest frontier, is an important driver of land use change (Foley et al., 2005), especially in Africa (Curtis et al., 2018). Ensuring that biodiversity conservation goals can be met without exacerbating poverty of these often already politically marginalized people (Fisher & Christopher, 2007), therefore requires high-quality information on the distribution of populations in these frontier regions. Unfortunately, the location and population density of forest-proximate people is poorly known and often badly captured by existing datasets including national gazetteers (Newton et al., 2020; Thomson et al., 2022) meaning many people are essentially invisible to conservation policy and practice (Poudyal et al 2016).

Understanding the distribution of human populations is needed to ensure conservation policy and practice is appropriately participatory and does not harm local people. A major expansion of area-based conservation measures is underway, following the agreement by nearly 200 countries to conserve 30% of the earth for nature by 2030 (CBD, 2022). Policymakers need to consider where people already live to reduce the opportunity costs, and potential conflict, associated with protected area establishment (Adams et al., 2010). The importance of genuine consultation with local people during the process of establishing new protected areas is recognized (Pimbert & Petty, 1997; Dawson et al., 2017), and requires information on the distribution of settlements.

Conservation researchers also need data on the density and distribution of human populations for a variety of purposes. This includes the selection of appropriate controls for exploring the impact of area-based conservation interventions on conservation outcomes (e.g. species populations; Wauchope et al., 2022, or tree cover loss; Desbureaux, 2021), or extrapolation from small-scale studies about the opportunity costs of conservation to the landscape scale, whether these costs arise through restrictions on agricultural expansion (Poudyal et al., 2018), or wildlife-human interactions (Drake et al., 2020).

A suite of open access global-scale datasets provide information on the distribution of human settlements. These are for example the World Settlement Footprint 2015 (Marconcini et al., 2020), the Open Buildings dataset (Sirko et al., 2021) or the WorldPop population density map (Tatem, 2017). However, several factors can limit the detectability of people living in forest-frontier regions including the often small size of settlements and potentially the lack of contrast between building materials and the surrounding environment (Ji et al., 2020; Van Den Hoek & Friedrich, 2021). There are few, if any, studies which explicitly test how well these datasets perform at detecting the distribution of forest-proximate people.

Detailed manually-derived data on the distribution of hamlets, villages and towns in the north-east of Madagascar (see Schüßler et al., 2020) offer an opportunity to test the reliability of global-scale remote-sensed datasets at capturing the density and distribution of human populations in a frontier area of great importance for biodiversity conservation. Madagascar’s highly threatened biodiversity is mostly located in the rapidly shrinking forests across the island (Ralimanana et al., 2022; Vieilledent et al., 2018), with shifting cultivation representing a major driver of ongoing forest loss (Curtis et al., 2018, though see McConnell and Kull, 2014). Sixty-five percent of the Malagasy population live in rural areas (UNDP, 2016), where food insecurity is particularly rife (Harvey et al., 2014; Thompson et al., 2023). Nationally-available datasets of the distribution of rural populations are largely absent (Ihantamalala et al., 2020). The country has undergone a substantial expansion of its protected area network (Gardner et al., 2018) which has sometimes negatively affected poor and already marginalized households living in forested areas (Jones et al., 2021; Rakotonarivo et al., 2017). However, previous attempts to compensate the local costs of conservation have failed (Brimont et al., 2015; Hockley et al., 2018; Poudyal et al., 2018), partly because of the limited available information on the location of forest-proximate people (Poudyal et al., 2016).

We evaluate the suitability of three global-scale remote-sensed datasets in detecting settlements and the density of forest-proximate people in north-eastern Madagascar. By comparing these datasets with a fine-scale manually-derived dataset (Schüßler et al., 2020), we explore the performance of the World Settlement Footprint and the Open Buildings Dataset at capturing the location of settlements, and WorldPop at capturing the density of human populations across the landscape. We discuss the factors associated with the detectability of forest-proximate people and the implications of these results for conservation policy, practice and research.

## Methods

### Study region

North-eastern Madagascar is characterized by the largest remaining rainforests of the island and a mosaic of regenerating fallows and farmland (Zähringer et al., 2015). While there are larger towns and cities along the more accessible coastline, inland there are scattered villages and hamlets. The main agriculture is shifting cultivation of rice, maize and manioc, supplemented with agroforestry production of cash crops such as vanilla, cloves or coffee (Llopis et al., 2019; Zähringer et al., 2015; Schüßler et al., 2020). We focus specifically on the Analanjirofo region which at 22,000 km² is of an equivalent size to El Salvador, Israel or Slovenia and which contains three major protected areas: Makira Natural Park, Mananara Nord National Park and Ambatovaky Special Reserve.

### Datasets

The World Settlement Footprint 2015 (hereafter WSF2015; Marconcini et al., 2020), indicates the presence or absence of human settlements in each grid cell of a raster at 10 m resolution. The Open Buildings Data (Sirko et al., 2021), provides the location of single buildings as shapefile data for the African continent. WorldPop (Tatem, 2017), is widely used raster data representing estimated population density at a resolution of 1 km. Details of the datasets, including the timestamp for data acquisition, is given in Table S1.

The fine-scale, manually-derived dataset was produced by Schüßler et al. (2020) to investigate the drivers of forest change in the Analanjirofo region. Schüßler et al. (2020) digitized the location of all villages detected visually from high-resolution images in Google Earth. The area was subdivided into seven parts and carefully scanned along north-south running transects for indications of human housing. The best quality and most cloud-free images were selected ranging in time between 2012 and 2018 (Table S1; Schüßler et al., 2020). Every settlement was marked as a point in its center and accompanied with its size (i.e., number of houses) in 8 size classes, ranging from 5-50 to 51-100 etc. to >350 houses per settlement. Many people in the study region have temporary houses in their agricultural fields but which are not permanent dwellings.

Settlements with 1-4 houses were therefore not recorded. This resulted in a total number of 3,136 settlements of which 59% consist of less than 50 houses, and a further 38% between 50 and 150 houses. Larger villages and cities made up the remaining 3% of all settlements of the area. While of course errors can be made in any dataset, for this analysis we treat this data as a measure of truth against which to compare the global datasets.

### Treatment of data

The three global-scale remote-sensed and the fine-scale manually-derived reference dataset are of different resolutions and of different data types (i.e., 10 m vs. 1 km, raster vs. shapefile). To explore the accuracy of the datasets at detecting settlements, we modified the data as follows. For comparability with WSF2015 (1x1 km raster data), we converted the fine-scale data to the same grid cell size, and recorded whether a grid cell size of 1km² contained a manually-derived settlement or not. The Open Buildings dataset was first converted from single buildings to settlements by creating 100 m buffers around all buildings and dissolving them to settlements (excluding groupings of fewer than 5 households as these had been excluded when preparing the manually-derived dataset). This buffer size was selected based on visual interpretation of satellite images overlaid with the Open Buildings dataset; increased buffer sizes of 200 or 300m did not change the results. Overlap of settlement locations between Open Buildings and our fine-scale manually derived dataset could then be measured and analyzed, as points of the latter dataset were located in village centers, warranting overlap with Open Buildings buffer polygons. For comparability with the WorldPop population density map, we calculated a kernel density map (Silverman, 2018) based on the number of houses per settlement in the fine-scale manually-derived dataset (and an estimate of 5.5 persons per household from Poudyal et al., 2018). These two population density datasets were then log-transformed and z-standardized to achieve a normal distribution and to increase comparability among datasets. Due to artefacts in the WorldPop data (indicated with strong outliers in the dataset and sharp edges in Figure 4), we removed outlier raster cells beforehand based on the 1.5*interquartile range rule (Yang et al., 2019). For comparability and illustrative purposes, we also produced a population density map from the Open Buildings dataset (again using an average estimate of 5.5 persons per household from Poudyal et al., 2018). We do not test this dataset against the data from the manually-derived dataset as we do not necessarily believe that for population density, the manually-derived data (based on point locations for settlements of varied size) is inherently better.

### Analysis

We used generalized linear models with a binary response variable (i.e., settlement detected or not) to compare both the WSF2015 and the Open Buildings datasets with the manually-derived fine-scale dataset. We used a linear model with the deviance between the population density estimates per grid cell as response variable for the comparison of WorldPop and the kernel density map from the manually-derived fine-scale data. To explain variation in detectability and density estimates of the three datasets we ran our models with six explanatory variables (described in Schüßler et al., 2020): (1) the density of forest per km² (in 2018) as a measure of proximity to the forest frontier (distance to the forest edge could not be used, as it does not distinguish between small forest fragments and the major forest blocks); (2) distance to major pathways of the study area (which represents accessibility); (3) distance to valley bottoms; (4) distance to major rivers (as preferred places for settlement formation); (5) the elevation and (6) the slope. Distance measures were calculated as Euclidean distances given in m. All explanatory variables were z-standardized to account for high variability in the scale of values. Correlations among explanatory variables were r < 0.65. Odds Ratios are presented for generalized linear models and standardized coefficients for the linear model. A further multi-model inference approach is detailed in the supplementary methods below.

## Results

### The accuracy of global-scale remote-sensed datasets at detecting settlements

Visual comparison of the two global-scale remote-sensed datasets with our fine-scale, manually-derived data suggests that the Open Buildings dataset detects a higher proportion of the settlements in our study area than the World Settlement Footprint data (WSF2015; Figure 1). The Open Buildings dataset detected a much higher proportion of the settlements than did the WSF2015 (94% or 2,914 of the settlements, compared to 15% or 484, Figure 2). WSF2015, performed particularly badly at identifying settlements of between 5 and 100 houses: 79% (2,477) were not detected while the Open Buildings dataset missed only 10% (177) of these small settlements (Figure 2).

**Figure 1:**
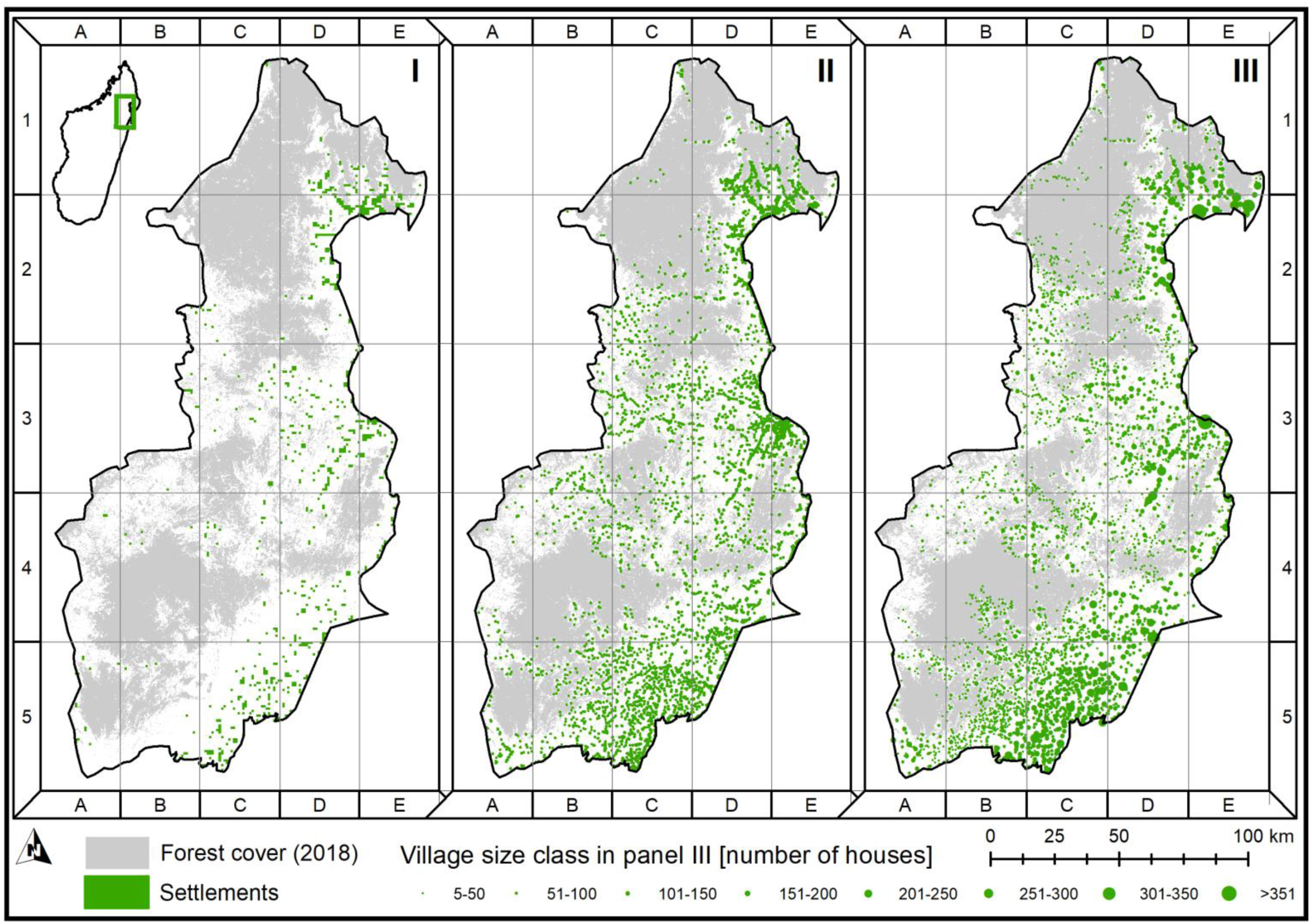
Distribution of settlements in the Analanjirofo region of north-eastern Madagascar as detected by I) World Settlement Footprint (WSF2015; raster data, 10 m resolution), II) Open Buildings (shape file of individual buildings) and III) our fine-scale manual-derived dataset (shape file with points of settlement locations).

**Figure 2:**
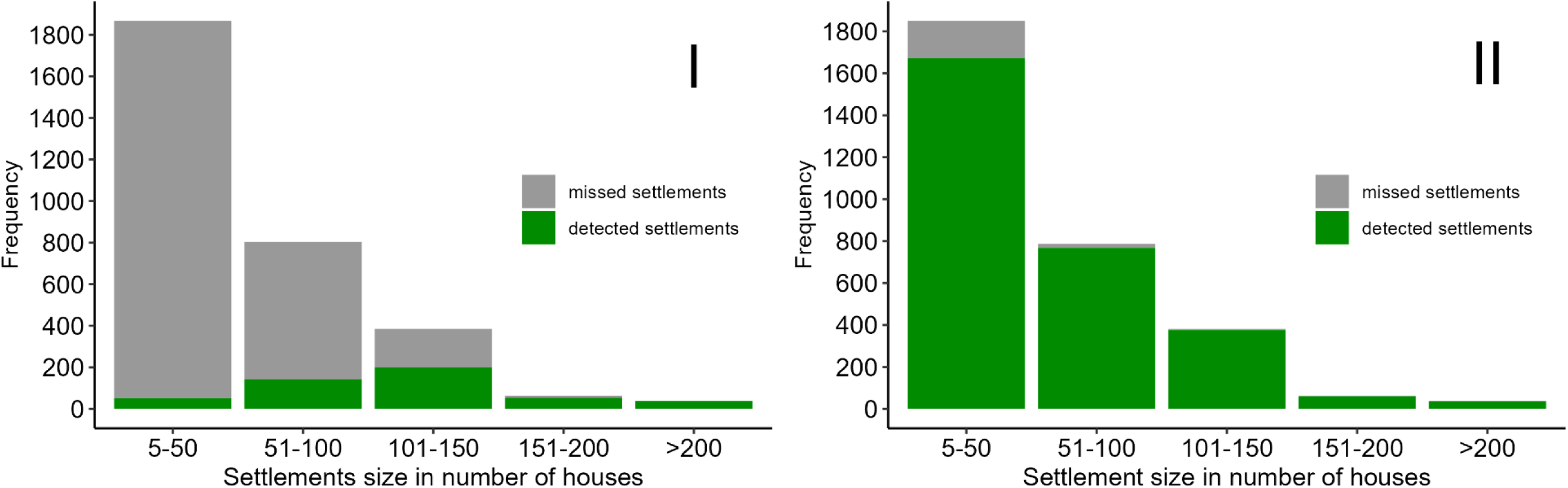
Number of settlements detected in different size categories by (I) World Settlement Footprint 2015 and (II) Open Buildings compared to the fine-scale manually derived dataset.

While WSF2015 clearly missed a lot of settlements across the landscape, the dataset was particularly likely to miss settlements in areas with high forest density, which were far from paths, at high elevation and further from valley bottoms (Figure 3I). Distance to major rivers did not contribute to the detectability of settlements and this variable was not included in the best performing model (Table S2 in SI). Given the higher agreement of the Open Buildings dataset with our manually-derived dataset in general (Figure 2), it is unsurprising that fewer variables significantly predict the detection of settlements by this dataset. However lower detection was similarly associated with areas of higher forest and higher elevations (Figure 3II). All other variables contributed to a lesser extent and were insignificant (Figure 3II; Table S2).

**Figure 3:**
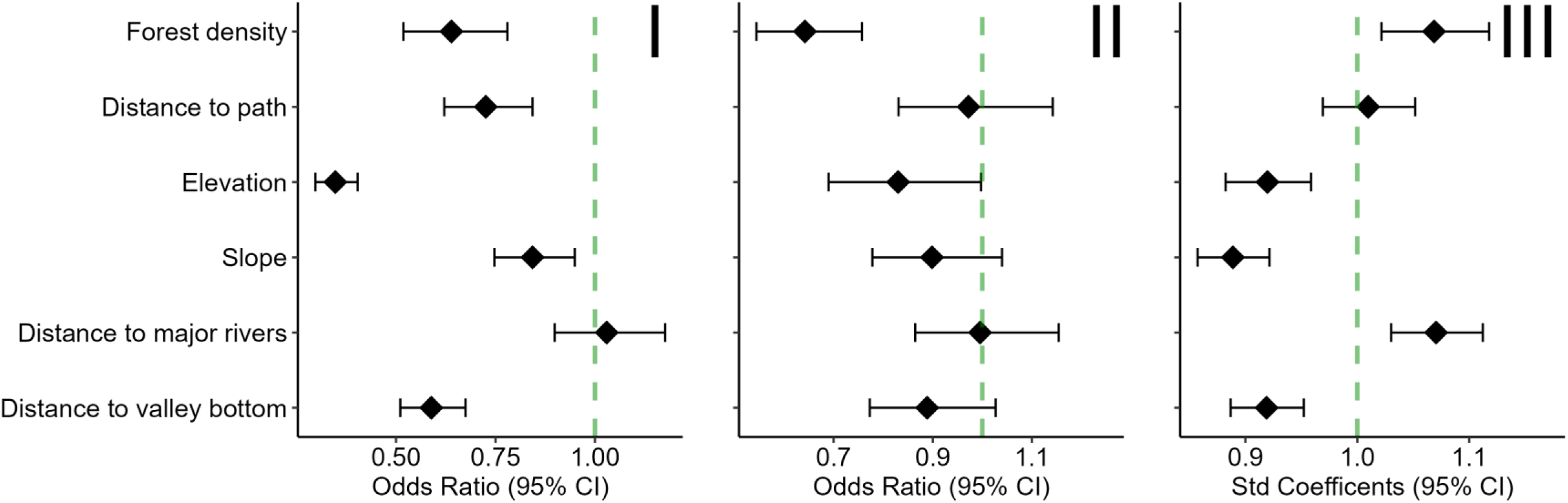
Effect sizes of the seven explanatory variables in explaining detection of settlements by the global-scale data (I, II) and the deviance in population density estimates (III). I: Odds Ratios of World Settlement Footprint 2015, II: Odds Ratios of Open Buildings dataset, II: Standardized coefficients of WorldPop. Calculation of Odds Ratios and standardized coefficients is based on global models containing all 7 explanatory variables (multi-model inference in Table S2). Note: Significant parameters are those not overlapping with the dashed line of no effect; values oriented in negative direction from the dashed line indicate lower detectability based on an increase in this parameter while values in positive direction from the dashed line show parameters that contributed to higher detectability of settlements based on the increase of this parameter.

### The accuracy of global-scale remote-sensed datasets at characterizing the variation in population density across the landscape

We were able to produce density estimates across the landscape from the Open Buildings dataset (Figure 4II) and the fine-scale manually-derived dataset (Figure 4III) and compared the latter to the density raster provided by WorldPop (Figure 4I). WSF2015 could not be easily converted into a comparable population density as it is presence/absence of settlements in a raster so it is not included in Figure 4. Apparent inaccuracies with sharp lines were found for many places in the WorldPop dataset (Figure 4I, cells C3 and C5). The density map estimates from the Open Buildings dataset displayed a pattern comparable to our manually derived dataset (Figure 4II, III). We did not model this relationship, as we produced the population density map ourselves, which may not directly allow to judge about the original dataset and the comparison of the original data was already conducted above (Figure 1-3).

**Figure 4:**
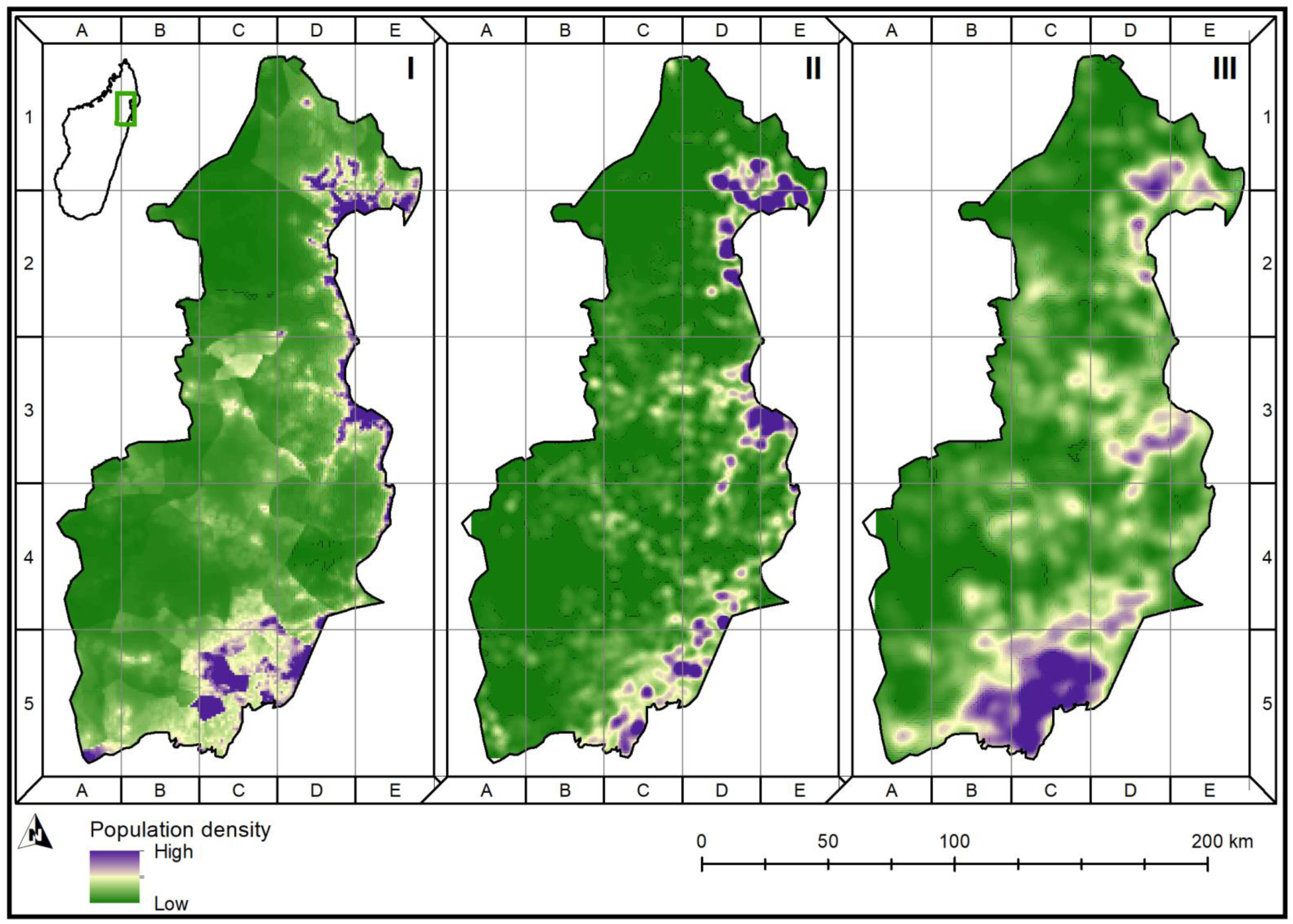
Population density in north-eastern Madagascar at 1 km resolution as derived from WorldPop (I), Open Buildings (II) and the fine-scale manually-derived dataset (III).

WorldPop was particularly poor at characterizing the population density in areas of lower forest density, at higher elevations, on steeper slopes and further away from valley bottoms (Figure 3III). Correct detection was higher in areas closer to the main rivers of the region. The best model did not include distance to pathways, but all other explanatory variables (Table S2) and explained only 3.4% of the variation. Although a high population density along the coast was found by WorldPop, it failed to estimate population density correctly further inland (Figure 4I, III).

## Discussion

### Which dataset performed best?

In many less developed countries, maps of human settlements are unavailable or inaccurate (Ihantamalala et al., 2020). Remote-sensed global datasets can help fill gaps, however, for such datasets to be used to inform conservation practice, policy or research, they need to accurately capture the location and density of human populations. Our results show that for detecting the distribution of human populations in a forest frontier region of Madagascar, the available remote-sensed datasets are of variable quality.

The Open Buildings dataset was particular promising in detecting both the location, and potentially density, of forest-proximate people. The Open Buildings dataset detected a much higher proportion of settlements in our manually-derived dataset than the World Settlement Footprint. This was especially true for the smallest settlements where Open Buildings dataset detected 90% compared to just 21% for World Settlement Footprint. Both datasets were particularly likely to miss settlements in more remote, forested areas and it is important to note that even the Open Buildings dataset missed 7% (177) of them.

While WorldPop is very widely used as an estimate of the distribution of human population density (at 1km resolution, Tatem, 2017), we found it performed rather poorly in our highly-forested study region. It further included sharp edges; an artefact from the processing of underlying datasets. Although the coastal regions were correctly shown to have high population densities, inland areas were inaccurate in both the visual interpretation (Figure 4) and the linear model explaining the deviations (Figure 3III). We did not formally test the performance of the density layer derived from the Open Buildings dataset. However, as this is based on detection of single buildings (rather than a classification of the approximate size of settlement as in our manually-derived dataset; Schüßler et al., 2020), this, when combined with a regionally-appropriate estimate of household size, may capture human population density most accurately (see Figure 4II).

### Limitations

There are a few limitations to our study. While we treat the manually-derived dataset as representing the truth for the purpose of this analysis, there will inevitably be some inaccuracies in this dataset. Firstly there is some discrepancy in the timestamp associated with the various datasets. As settlements can appear (and less commonly disappear) relatively rapidly in frontier regions (Foley et al., 2005), lack of accuracy detected in the remote-sense datasets may relate to inconsistencies in which settlements were present at a moment in time, rather than ability of certain approaches to detect settlements of a given size. However, all datasets used in this study (with the exception of the Open Buildings dataset for which not date is defined) come from a relatively short period (see Table S1), so that we do not consider this to be a major problem. Another limitation with the manually-derived dataset is that it does not include individual isolated homesteads. While many isolated buildings represent temporary shelters for use when tending crops at certain times of the year, others will represent permanent dwellings. By not recording settlements with fewer than 5 houses our dataset risks missing some of the most isolated people in the landscape.

Our results may further only be applicable to settlement detectability in humid tropical regions. Similar analysis would be needed to explore the extent to which the findings are transferable to arid regions. This is because single houses (as in the Open Buildings dataset) and settlements (as in the WSF2015 dataset) likely contrast better with the surrounding evergreen vegetation than might be true in more arid environments.

### Implications for forest-proximate communities

There are at least 1 billion forest-proximate people in tropical countries (Newton et al., 2020). Such people are often marginalized politically and economically (e.g., Levers et al., 2021) and conservation has often proceeded without appropriate involvement of these key stakeholders (Adams et al., 2010). The lack of high-quality information about the distribution of people at the forest frontier has effectively made such people invisible. The aim of the study was to highlight the potential for remote-sensed datasets to make them more visible: easier to consider in conservation policy, easier to reach for consultation or involvement in conservation practice, and easier to take into account in conservation research. Our findings confirm the value of such datasets, and suggest that the Open Buildings dataset has particular value (at least in a forest frontier region of north-eastern Madagascar). However even the best dataset struggles to capture settlements in the most forested areas. If this isn’t addressed, there is a risk of these people continuing to be overlooked.

## Acknowledgement

This work was funded by the UK Government CLARE programme (Climate Resilience and Adaptation) and the International Development Research Centre, Ottawa, Canada through their support for the Forest4Climate&People project. We thank Neal Hockley and Bruno Ramamonjisoa for support during this project and Ute Radespiel for supervision during the generation of the manually-derived fine-scale dataset. We further thank Dr. Enrico Di Minin for valuable comments on an earlier version of the manuscript. The authors declare not conflict of interests.

## Author contribution statement

JPGJ and DS conceived of the ideas for this manuscript. MR and DS collated and analyzed the data and lead the manuscript writing. All authors contributed to previous drafts of the manuscript, and approved the final version for publication.

## Data availability statement

The global datasets (WorldPop, WSF2015, Open Buildings) used in this manuscript are freely available. The fine-scale manually-derived dataset can be shared upon request to the corresponding author.

## Further statements

The authors declare no conflict of interests.

## Supplementary information

### Supplementary methods

The spatial data was freely available from Tatem (2017), Sirko et al. (2021) and Marconcini et al. (2020) and provided by Schüßler et al. (2020). Spatial data was prepared in ArcGIS 10.6 (ArcGIS Desktop 10.6.1, ESRI, Redlands, USA) and QGIS 3.10 (QGIS Development Team, 2020) and later analyses were conducted in R v4.4.2 and RStudio v2022.12.0.353 (R Core Team, 2022; Posit Team, 2022). We checked for collinearity of the seven explanatory variables beforehand using the “psych” package v2.2.9 in R (Revelle, 2022) and all correlations were below r = 0.65.

Supplementary Table S1. provides details on the used datasets and their specifications. Using a multi-model inference approach (Burnham & Anderson, 2002) for each comparison (i.e., WSF2015 and Open Buildings with the manually-derived dataset, WorldPop with the kernel density estimates from the manually-derived dataset), we fitted a global model first with all six explanatory variables (i.e., forest density, distance to major pathways, distance to main rivers, distance to valley bottoms, slope and elevation). Models containing all possible combinations of variables were then calculated using the “MuMIn” package v1.47.1 (Barton, 2022). Candidate models based on the lowest AIC values and those deviating less than 2 units from the best model (ΔAIC<2) are reported in this study (Burnham & Anderson, 2002). Akaike weights (i.e., conditional probability of a certain model) were calculated based on all possible models (Wagenmakers & Farrell, 2004). Model results from the generalized linear models are presented as Odds Ratios calculated using the “effectsize” package v0.8.2 (Ben-Shachar et al., 2020) and visualized using “ggplot2” v.3.4.0 (Wickham, 2016) and “patchwork” v1.1.2 (Pedersen, 2020). For the linear model, standardized coefficients are presented.

### Supplementary results

Forest density and elevation were the only variables present in all candidate models as a predictor of the detectability of settlements or correct estimation of population densities (Table S2); the other variables were less often involved. The best model on the detectability of settlements of WSF2015 involved elevation, forest density, distance to valley bottoms, distance to pathways and slope as explanatory variables and yielded the highest Akaike weight with 0.682. The other candidate model was of subordinate quality (Table S2).

For Open Buildings, the best model involved forest density, elevation, distance to valley bottoms and slope as explanatory variables. However, multiple further candidate models with only slightly lower Akaike weights involved also all other variables at some point. The best model concerning correct estimation of population density by WorldPop included all explanatory variables, except distance to pathways and yielded an Akaike weight of 0.693, markedly higher compared to the second best and only other candidate model (Table S2).

**Supplementary Table S1:** Details on the remote-sensed datasets used in this study.

| Dataset | Data type | Resolution | Acquisition date | Description |
| --- | --- | --- | --- | --- |
| WorldPop<br>(Tatem,<br>2017) | Raster | 30 arc-seconds<br>(approx. 1 km at<br>the equator) | 2015 | This is an estimated population density per raster grid cell. The units are the number of people per square kilometer. |
| WSF2015<br>(Marconcini<br>et al., 2020) | Raster | 10 m | 2014 – 2015 | This dataset shows the presence or absence of human settlements in each raster grid cell in 2015. It has been derived by jointly exploiting multi-temporal Sentinel-1 radar and Landsat-8 optical satellite imagery. Its quality has been validated against 900,000 sample localities labelled by crowdsourcing photointerpretation of very high-resolution Google Earth Imagery. The temporal resolution from Landsat-8 and Sentinel-1 are 16 days and 6 days, respectively. |
| Open<br>Buildings<br>(Sirko et al.,<br>2021) | Polygone-<br>shapefile | - | Not defined, different timestamps | This dataset contains 516 million buildings across the African continent. It has been generated by using 50 cm satellite imagery from Maxar Technologies and CNES Airbus. |
| Manually-derived fine-scale dataset<br>(Schüßler et<br>al., 2020) | Point-<br>shapefile | - | Different timestamps, split by region:<br>• Makira west: 09/07/2014<br>• Masoala west: 09/07/2014<br>• Antainambalana river: 09/11/2017<br>• Makira north: 01/06/2013<br>• Mananara west: 09/07/2014<br>• Makira south: 15/05/2012<br>• Makira southwest: 07/07/2013<br>• Mananara west: 12/09/2015<br>• Southwestern edge of region: 15/06/2015<br>• Southeastern edge of study region: 31/12/2016<br>• Masoala northwest: 12/09/2015 | This dataset was manually extracted from high resolution Google Earth imagery. Settlements were digitized as point localities in their middle and categorised in seven classes according to the number of visible house:<br>• 5-50 houses: 59% of settlements<br>• 50-150 houses: 38% of settlements<br>• >150 houses: 3% of settlements |
|  |  |  | • Mananara Nord National Park: 06/06/2018<br>• Makira southeast: 09/07/2014<br>• Makira northeast: 09/11/2017<br>• Soanierana-Ivongo: 05/02/2017<br>• Ambatovaky Special Reserve: 12/09/2015<br>• Marotandrano Special Reserve: 22/01/2013<br>• Southern edge highlands: 15/06/2015<br>• Anove river highlands: 12/09/2015 |  |

**Supplementary Table S2:** Multi-model comparison to explain differences in village detectability (WSF2015 and Open Buildings via GLMs) and deviance in population density estimates (WorldPop via LM). Candidate models with ΔAIC<2.0 are shown here with the best model given in bold. Akaike weights were calculated on all possible models and indicate how well this model performs in comparison to the other candidate models. Empty cells indicate that these parameters were not included in the respective model. Abbreviations: Int. = intercept, elev = elevation, for. dens. = forest density, AIC = Akaike Information Criterion.

| | Int. | Elev. | For. dens. | Major river | Valley bottom | Path | Slope | $\Delta AIC$ | Akaike weight |
| --- | --- | --- | --- | --- | --- | --- | --- | --- | --- |
| WSF2015 | **-3.767** | **-1.247** | **-0.628** |  | **-0.682** | **-0.414** | **-0.174** | **0.00** | **0.682** |
|  | -3.760 | -1.252 | -0.633 | 0.040 | -0.686 | -0.416 | -0.172 | 1.82 | 0.274 |
| Open Buildings | **2.382** | **-0.221** | **-0.581** |  | **-0.153** |  | **-0.106** | **0.00** | **0.128** |
|  | 2.367 | -0.237 | -0.547 |  | -0.163 |  |  | 0.08 | 0.123 |
|  | 2.443 | -0.242 | -0.614 |  |  |  | -0.116 | 0.61 | 0.095 |
|  | 2.432 | -0.260 | -0.571 |  |  |  |  | 1.08 | 0.075 |
|  | 2.377 | -0.221 | -0.574 |  | -0.154 | -0.039 | -0.107 | 1.85 | 0.051 |
|  | 2.363 | -0.236 | -0.561 |  | -0.164 | 0,038 |  | 1.93 | 0.049 |
|  | 2.381 | -0.217 | -0.571 | -0.016 | -0.152 |  | -0.109 | 1.97 | 0.048 |
| WorldPop | **-0.328** | **-0.054** | **-0.054** | **-0.052** | **-0.060** |  | **-0.065** | **0.00** | **0.693** |
|  | -0.327 | -0.054 | -0.051 | -0.051 | -0.060 | 0.006 | -0.065 | 1.76 | 0.283 |

